# Proteomic Analysis in Alzheimer’s Disease with Psychosis Reveals a shared AD proteomic landscape and a separate molecular signature of Postsynaptic Density Disruption

**DOI:** 10.1101/2025.11.26.690872

**Authors:** T.S. Ku, S.J. Mullett, Z. Sui, L. Zeng, Y. Ding, A.K. Yocum, M. MacDonald, S.L. Gelhaus, J Kofler, R.A. Sweet

**Affiliations:** Dept. Psychiatry, Univ. of Pittsburgh; Dept. Biostatistics and Health Data Science, Univ. of Pittsburgh; Dept. Pharmacology and Chemical Biology, Univ., Pittsburgh; Dept. Pathology, Univ. of Pittsburgh; Dept. Neurology, Univ. of Pittsburgh

**Author notes:** Corresponding author: Robert A. Sweet, MD, 3811 O’Hara St., Pittsburgh, PA, 15213.

## Abstract

**Background and Hypothesis:** Alzheimer’s disease with psychosis (AD+P) is a subgroup of AD patients with more rapid cognitive deterioration. While our previous study showed that AD+P is associated with loss of prefrontal cortex postsynaptic density (PSD) proteins, identifying proteins in the broader cellular environment that influence PSD loss addresses a critical knowledge gap about synaptic dysfunction mechanisms in early disease stages.

**Study Design:** We conducted a proteomic analysis comparing prefrontal grey matter cortex tissue homogenates from elderly normal controls (n=18), individuals with AD+P (n=61), and individuals with AD-P (n=48).

**Study Results:** AD+P showed the most pronounced alterations relative to controls (178 proteins with q<0.05), although alterations in AD-P and AD+P relative to controls were highly similar (R²=0.965, p<0.001). Weighted-gene correlation network analysis (WGCNA) identified four modules significantly associated with disease status comparing AD subjects to controls, but none differed significantly between AD+P and AD-P. We identified 15 proteins significantly correlated with PSD yield across all samples, including ENPP6, linked to AD+P by GWAS. Additionally, PSD yield-associated proteins showed minimal overlap with altered AD proteins (1 of 137). Proteins negatively correlated with PSD yield were enriched for presynaptic endocytosis, whereas the one WGCNA module significantly correlated with PSD yield was enriched for inflammatory terms.

**Conclusions:** Our findings suggest a model in which AD+P arises from the combination of quantitative alterations within a shared AD proteome profile and a superimposed set of protein alterations correlated with PSD yield that confer distinct mechanisms of synaptic vulnerability and psychosis risk.

## Introduction

Alzheimer’s disease (AD) stands as the most prevalent neurodegenerative disorder and the leading cause of dementia worldwide^1^. The pathological changes, characterized by amyloid-beta plaques and neurofibrillary tangles, lead to neuronal dysfunction, synaptic loss, and ultimately, brain atrophy. AD manifests clinically as progressive memory impairment, cognitive decline, and deterioration in the ability to perform daily activities (Reviewed in^2^). Critically, synapse loss represents the earliest and the strongest correlate of cognitive decline in AD (Reviewed in^3^).

Psychotic symptoms, including delusions and hallucinations, affect approximately 40%-60% of individuals with AD^4^. The occurrence of psychosis in AD (Alzheimer’s disease with psychosis, AD+P) is a marker for a subgroup of AD patients with more rapid cognitive deterioration^5^. However, only a minority of the risk for the more severe AD+P phenotype is explained by the burden of AD and comorbid pathologies^6^. Instead, AD+P has been found to be associated with loss of prefrontal cortex postsynaptic density (PSD) proteins relative to both AD-P and elderly control subjects^7^. While genome-wide association studies and transcriptomic analyses^8,9^ have provided insights into additional molecular risks for the AD+P subgroup, the specific proteomic landscape and its relationship to PSD integrity remains poorly understood.

The postsynaptic density is a protein-rich structure at excitatory synapses that is essential for synaptic transmission and plasticity. It anchors glutamate receptors (NMDA and AMPA receptors) and links them to kinases and phosphatases through scaffolding proteins like PSD-95, enabling the regulation of synaptic strength through long-term potentiation (LTP)^10,11^. In AD, the PSD undergoes profound disruption, with PSD-95 levels decreased by approximately 19% compared to controls^12^. Even in mild cognitive impairment, often prodromal to AD, significant reductions in PSD-95 and associated proteins like NR2A and LRP1 are evident in the hippocampus^13^. Such widespread PSD compositional changes fundamentally compromise synaptic mechanisms essential for cognitive function^14^.

While previous studies have characterized PSD alterations in AD+P, upstream factors that initiate these disruptions remain poorly understood. Identifying proteins in the broader cellular environment that influence PSD loss addresses a critical knowledge gap about synaptic dysfunction mechanisms in early disease stages. To address this gap, we examined the prefrontal cortex gray matter proteome in individuals with early to moderate neuropathologic stages of AD and elderly controls in all of whom PSD abundance had been previously characterized. We found that AD+P was associated with a profile of gray matter proteome alterations relative to control subjects that was shared with AD-P, but of greater severity. In addition was a superimposed set of protein alterations that correlated with PSD yield.

## Methods

### Subjects & Sample Preparation

Frontal cortex gray matter from subjects with AD and from elderly cognitively normal control subjects (Supplementary Data 1) were obtained from the brain bank of the Alzheimer Disease Research Center (ADRC) at the University of Pittsburgh and from the Rush Alzheimer’s Disease Center, neuropathologically characterized, and processed to generate homogenate protein samples as previously described^7^. Control subjects were Braak stages 0–2, whereas AD subjects were restricted to Braak stages 3–5; end-stage cases, as defined by a Braak stage of 6, were excluded. Psychosis was defined as the presence of delusions or hallucinations at any clinical visit prior to death as previously described^15^.

10 µg total protein homogenate from each sample and pooled “trypsin digestion” controls were randomly assigned to four blocks for trypsin digestion on S-TRAP 96 well plates (Protifi) following manufacture protocols, and two blocks for desalting. Block randomization was stratified by and balanced for diagnosis groups (AD+P/AD-P/NC). Chi-square test and t-test showed that the design was balanced for diagnosis groups, sex, age and PMI (p>0.05). Within each block for digestion or desalting, run orders were generated randomly. Run orders for mass spectrometry were generated randomly across all samples. Additionally, 100 µg of pooled sample was digested on S-TRAP 96 Mini Columns (Protifi), fractionated with the Pierce™ High pH Reversed-Phase Peptide Fractionation Kit (Thermo Fisher) for the generation of a custom DIA library.

**Supplementary Figure 1.**
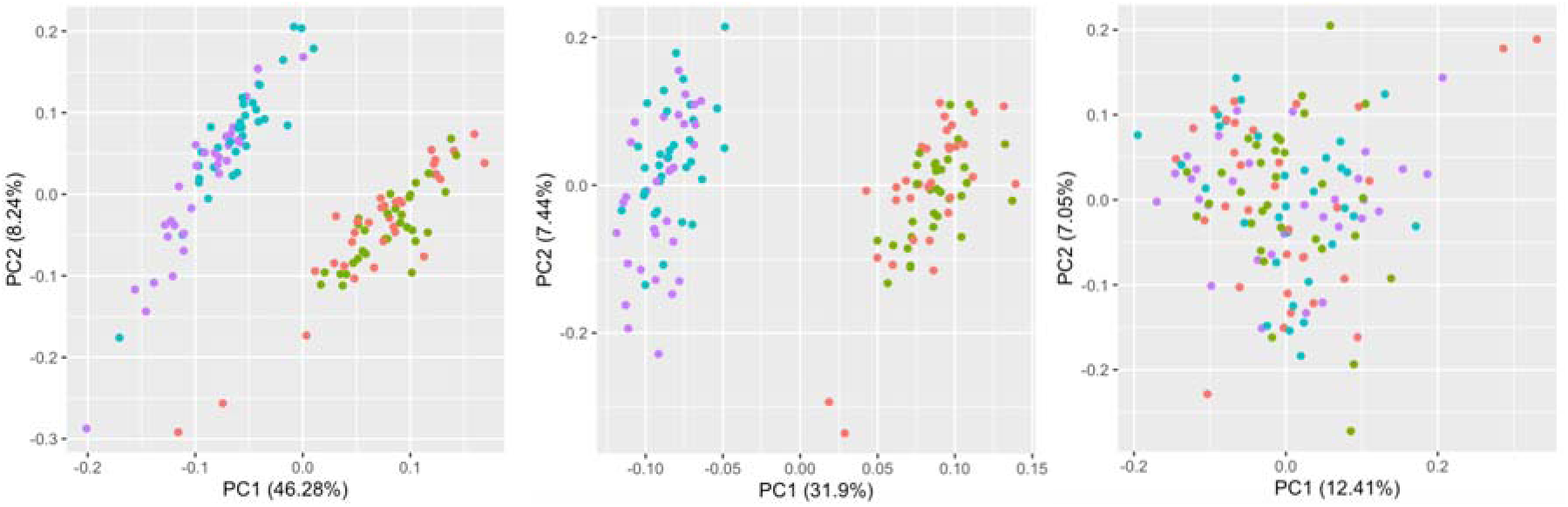
Effects of normalization on proteomics data assessed by principal component analysis (PCA). PCA plots of samples before normalization (left), after median normalization (middle), and after median plus block normalization (right). Each point represents an individual sample and colors indicate experimental blocks.

### DIA Data Analysis and Spectral Library Generation

Data-independent acquisition (DIA) data were analyzed using DIA-NN (v1.9.2). A project-specific spectral library was first generated with Data-dependent acquisition (DDA) analysis of the fractionated pooled sample described above, with peptide and protein identification guided by the UniProt human FASTA database (UP000005640, downloaded January 1, 2024; including decoys and common contaminants).

DIA-NN performed in silico spectral library generation using deep learning-based peptide prediction from FASTA digestion. Digestion parameters specified cleavage at lysine (K) and arginine (R) residues, allowing up to one missed cleavage, with peptide lengths ranging from 7 to 30 amino acids. Carbamidomethylation of cysteine was included as a fixed modification, and oxidation of methionine (UniMod:35) was permitted as a variable modification. Precursor ions were selected from m/z 500 to 1800, with charge states between +2 and +4, and fragment ions considered within the m/z range of 200 to 1800. Mass accuracy was fixed to 1.5e-5 (MS1) and 2e-5 (MS2), and only peaks with a correlation sum >2 were retained, while peaks with correlation <1 from the maximum were excluded. Retention time (RT) alignment was performed empirically, using a RT window of 2.44 minutes, ion mobility window of 0.041, and a peak width of 3.34.

The resulting spectral library contained 54,327 protein groups, 25,435 proteins, and over 6 million precursors, and was subsequently used for quantifying the full cohort of patient samples.

### MS Analysis

Mass spectrometry analysis was conducted on a Bruker NanoElute 2 UHPLC system coupled to a timsTOF Pro2 using CaptiveSpray nano-electrospray. Approximately 200 ng of peptide digest (2 µL) was loaded onto an Aurora Ultimate C18 column (1.7 µm, 25 cm x 75 µm, IonOpticks) equilibrated at 55°C and eluted at 300 nL/min over a 60-minute gradient. Mobile phase A consisted of 0.1% aqueous formic acid and mobile phase B was 0.1% formic acid in acetonitrile. A linear gradient of 2–35%B was applied for 60 min, followed by a 5 min wash at 95% B before column equilibration at the initial conditions of 2%B for 6 min. Each sample was analyzed once, without technical replicate injections.

The timsTOF Pro2 was set to DIA-PASEF scan mode with a scan range of 100 - 1700 m/z. The TIMS was set to a 100 ms ramp and accumulation time (100% duty cycle) with a ramp rate of 9.43 Hz. Cross sectional collision energy was set to a base of 1.60 1/K0 [V-s/cm2] at 59 eV and 0.60 1/ K0 [V-s/cm2] at 20 eV. Fragment ions were analyzed in 43 DIA-PASEF windows in a mass range of 252 – 1328 Da. An additional pooled “mass spec” control which combined 2µL of post-digestion peptides from every sample in the study were injected at regular intervals within the injection sequence to monitor instrument sensitivity and reproducibility.

### DIA Data Processing and Quantification

Patient samples were analyzed using the same version of DIA-NN (v1.9.2), utilizing the spectral library described above. Results were filtered at a 1% FDR for both precursor and protein levels. Protein inference was based on proteotypic peptides, and heuristic protein grouping was again used^16^. DIA-NN performed digestion using the same cleavage and modification parameters as in library generation, with the allowance of up to two variable modifications per peptide. For quantification, mass accuracy was fixed at 2e-5 for both MS1 and MS2, and calibration across runs resulted in a recommended MS1 accuracy of approximately 11.6 ppm. RT and ion mobility settings were refined to a 2.05-minute RT window, 0.0402 ion mobility window, and peak width of 3.59. Interference removal was disabled during this run.

### Statistical Analysis

Samples with more than 80% missing values across all peptides were removed. One sample met this criterion and was therefore excluded prior to subsequent analyses. A total of 127 biological samples remained: 48 with AD-P, 61 with AD+P, and 18 Normal Controls. We performed rigorous quality control and normalization on all quantified peptides. Peptides with missing rates >50% across all biological samples were removed. Additionally, peptides with fewer than three out of eight pooled control samples or with a coefficient of variation >100% in the pooled controls were removed. For each unique stripped peptide sequence, we retained the precursor with the smallest coefficient of variation among the pooled control samples. This filtering process resulted in 19,617 peptides for analysis.

We first applied median normalization to biological samples to make the median of each sample equal to the median of all samples. Peptide intensities were then log2-transformed, and the block effect from the digestion block was regressed for each peptide, with the residuals retained for downstream analysis (Supplementary Figure 1).

Next, we rolled up the peptides to protein-level using the inverse-CV-weighted average of scaled peptide values, where each scaled peptide value is the Z-score of the (log2-transformed) peptide abundance across samples. The same approach was used for targeted proteomics data in our previous work^15,17,18^. To investigate the relationship between protein abundance and psychosis diagnosis, for each protein, we fitted a multiple linear regression with protein abundance as the dependent variable, psychosis diagnosis as the independent variable, adjusting for sex, age, PMI, and APOE4 status. The regression analyses were conducted for psychosis diagnosis contrasts of AD+P vs. AD–P, AD+P vs. control, AD–P vs. control, and AD vs. control.

Then we applied Weighted Correlation Network Analysis (WGCNA)^19^ on all protein-level data to detect protein co-expression modules. A signed weighted correlation network was constructed using a soft-thresholding power of 4 to approximate scale-free topology. Modules were identified by hierarchical clustering with dynamic tree cutting using a minimum module size of 25 proteins, resulting in 16 distinct modules ranging in size from 50 to 1558 proteins. For each module, we obtained the module eigengene value and fitted a similar multivariate linear regression with module eigengene as the outcome, psychosis diagnosis as the predictor, adjusting for sex, age, PMI, and APOE4 status.

We also analyzed the correlation between protein abundance and PSD yield. We previously reported in these subjects that PSD yield (μg of PSD protein per μg of gray matter protein) was significantly reduced in AD+P relative to both AD-P and elderly cognitively normal subjects (NC: 0.526±0.201; AD-P: 0.513±0.241; AD+P: 0.421±0.192)^7^. Pearson’s correlation was then computed between each protein’s abundance and the PSD yield.

### Overrepresentation Analysis (ORA)

Overrepresentation analysis (ORA) was performed to identify biological pathways enriched among differentially abundant proteins. For each pairwise comparison (AD+P vs AD-P, AD+P vs NC, AD-P vs NC, and AD vs NC), proteins with q-value < 0.05 were selected as the input gene set for enrichment analysis.

We used the clusterProfiler R package (v4.14.6)^20,21^ to identify Gene Ontology (GO) Biological Process terms enriched among differentially abundant proteins (q < 0.05). UniProt IDs were mapped to Entrez Gene IDs via bitr() using org.Hs.eg.db (v3.20.0)^22^; unmapped proteins were excluded. The background universe included all successfully mapped proteins from the dataset. Pathways with q < 0.05 and q < 0.1 were considered significant. This slightly more permissive FDR threshold for pathway analyses (compared with protein-level differential abundance analyses) was applied to improve sensitivity for detecting coordinated biological processes. Comparisons with fewer than three significant proteins were excluded. Results were visualized with ggplot2 (v3.5.2) as dot plots showing the top 10 GO terms ranked by gene ratio.

### SynGO

Synaptic gene ontology (GO) enrichment analysis was performed using SynGO (release 1.3; www.syngoportal.org)^23^. Differentially expressed proteins (q < 0.05) from each of the three comparisons (AD vs. NC, AD+P vs. NC, and AD−P vs. NC) were uploaded separately to the SynGO web portal as input lists. The total detected proteome (n = 4,705) was used as the background reference set. A minimum of 3 proteins per GO term was required for inclusion in the analysis. Both biological process and cellular component ontologies were examined.

### Gene Set Enrichment Analysis (GSEA)

Gene set enrichment analysis (GSEA) was performed using the Broad Institute GSEAPreranked tool (GSEA v4.4.0)^24,25^ on proteins ranked by their correlation coefficients with PSD abundance. Proteins were ranked from highest positive to lowest negative correlation, and enrichment was assessed using the Gene Ontology Biological Process (GO BP) database with 1,000 gene set permutations. Gene sets with sizes between 15 and 500 genes were included, and enrichment significance was evaluated using normalized enrichment scores (NES) and false discovery rate (FDR) q-values.

## Results

### Protein Abundance Changes in AD with Psychosis

To characterize the molecular changes associated with Alzheimer’s disease with psychosis, we performed quantitative proteomic analysis on dorsolateral prefrontal cortex (DLPFC) samples from 127 individuals, including 48 AD-P, 61 AD+P, and 18 normal controls. Our analysis encompassed 4,705 proteins across all sample groups, providing a broad coverage of the brain proteome (Supplementary Data 2).

Differential expression analysis revealed that AD+P drives the most pronounced molecular alterations, with 180 proteins significantly altered (q < 0.05), while 55 proteins with q < 0.05 were identified comparing AD-P with control (Fig. 1A). Overlap analysis showed shared significant protein changes between AD groups and controls (Fig. 1B). Examining proteins significantly altered in AD+P relative to controls, we observed a high degree of correlation with the changes in AD-P relative to controls (Fig. 1C, left), but of greater magnitude in AD+P reflected by larger deviations from the identity line. In contrast, proteins that were significantly altered in both AD-P and AD+P relative to controls showed a strong directional consistency and close alignment with the identity line. (Fig. 1C, right). Together, these indicate that while both groups share a core pathological proteome signature, psychosis in AD is associated with amplified alterations in a subset of proteins. The most significantly altered core proteins (i.e. of those significantly altered in all AD versus control) included TAGLN, PRELP, LUM, CST3, BGN, and PKP2, which also emerged as top hits in the individual AD-P versus NC and AD versus NC comparisons (Fig. 1A). The comparison between AD+P and AD-P revealed proteins with only modest fold changes. Several proteins showed nominally significant differences (p < 0.05), with the most notable changes observed in CLPP (upregulated) and ITPK1 (downregulated).

**Figure 1.**
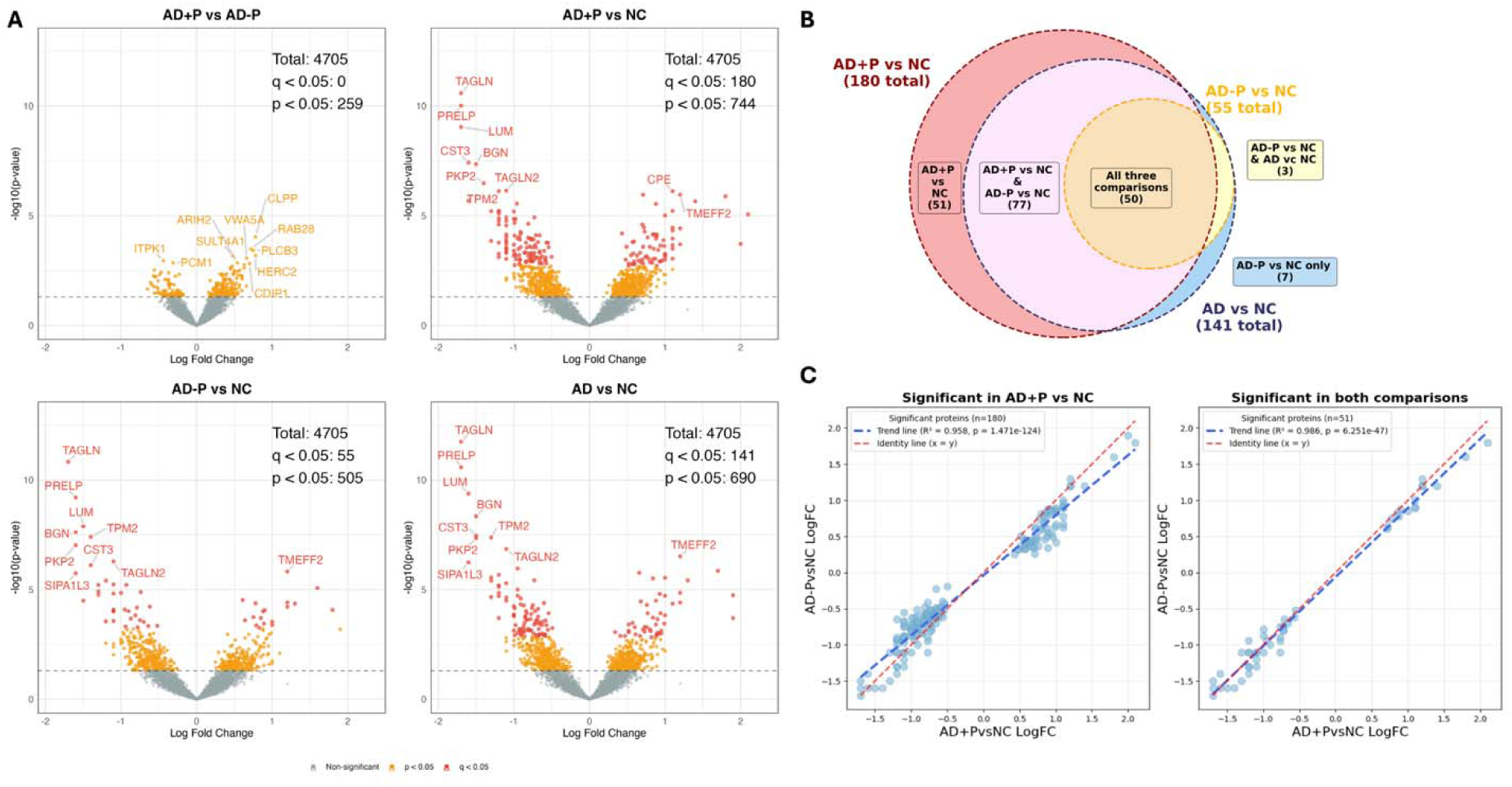
Differential Protein Expression across Alzheimer’s Disease Subgroups. **A.** Volcano plots displaying log2 fold change versus -log10(p-value), with proteins meeting q<0.05 highlighted in red and proteins with p<0.05 in orange. Top left: AD+P versus NC comparison. Top right: AD+P versus AD-P comparison. Bottom left: AD-P versus NC comparison. Bottom right: AD versus NC. **B.** Venn diagram illustrating overlap of significantly dysregulated proteins (q<0.05) among AD+P vs NC, AD-P vs NC, and AD vs NC comparisons. **C.** Correlation analysis comparing protein log₂ fold changes across Alzheimer’s disease subgroups. Left: Among proteins significantly altered in AD+P (q<0.05, n=180), the comparison of AD+P vs NC and AD-P vs NC reveals strong correlation (R²=0.958, p=1.471e-124). Deviation from the identity line indicates that proteins show overall larger magnitude changes in AD+P than in AD-P. Right: Proteins that are significantly altered relative to NC in both AD-P and AD+P are highly correlated with the magnitude of changes in each AD group close to identity (R^2^=0.986, p=6.251e-47, n=51 proteins significant in both comparisons with q<0.05). Together, these indicate substantial overlap in the nature of AD-related proteomic changes regardless of psychosis status, but of greater severity in AD+P.

### Pathways Enriched in Proteins Altered in AD+P

Functional analysis using Gene Ontology (GO) terms revealed striking similarities in biological processes across comparison groups (Fig. 2, Supplementary Data 3-5). All three comparisons (AD+P vs NC, AD-P vs NC, and AD vs NC) showed remarkably consistent patterns, with cell adhesion, supramolecular fiber organization, circulatory system development, extracellular matrix organization, extracellular structure organization, and collagen fibril organization being consistently enriched across groups. This pattern suggests that the core molecular changes in Alzheimer’s disease are largely the same regardless of whether patients present with or without psychotic symptoms. To further examine synaptic alterations, SynGO enrichment analysis was performed on differentially expressed proteins from each comparison group (Supplementary Figure 2). Of those proteins, 20/141 genes in AD vs NC, 29/180 genes in AD+P vs NC, and 7/55 genes in AD−P vs NC mapped to SynGO-annotated synaptic genes. These mapped proteins spanned multiple synaptic compartments and processes; however, no significant convergence on specific synaptic functions or pathways was observed across comparisons.

**Figure 2.**
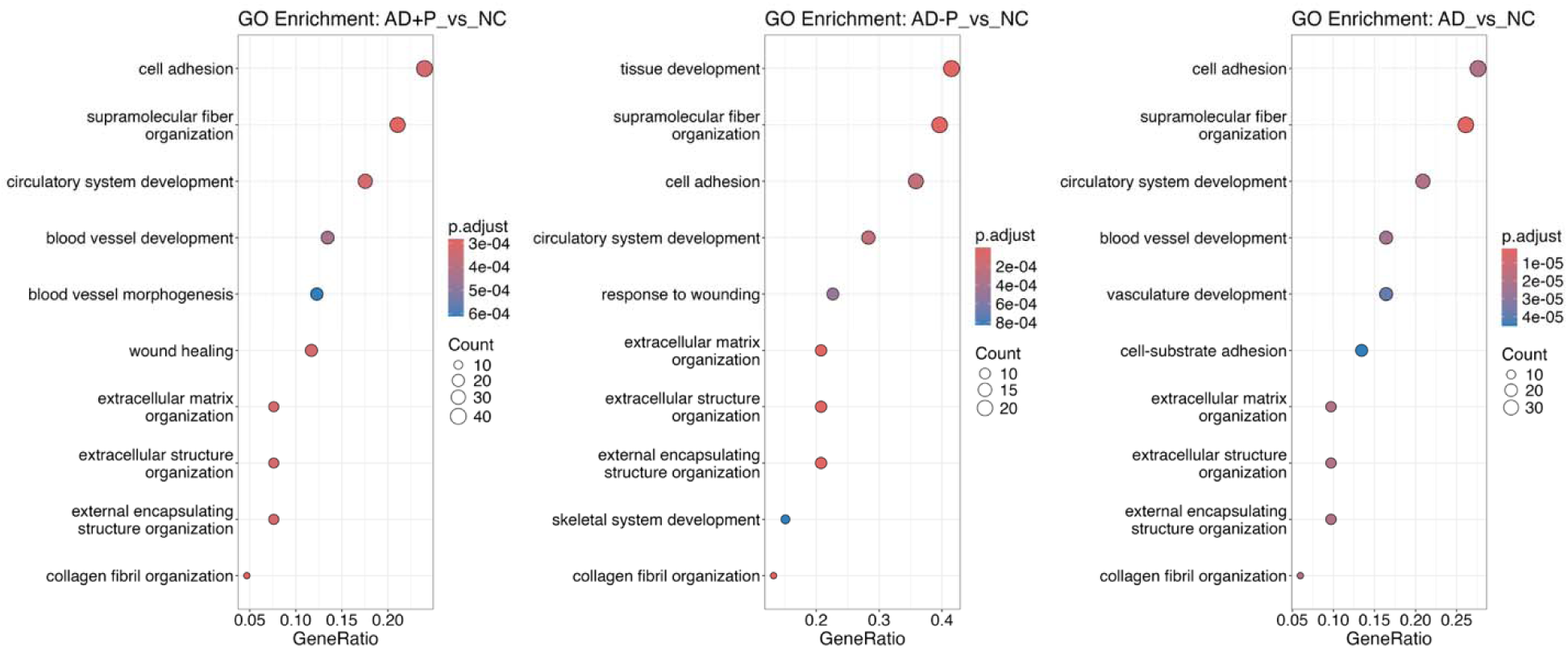
Over-Representation Analysis of Differentially Expressed Proteins. Gene Ontology (GO) enrichment analysis was performed on differentially expressed proteins (q < 0.05) against the background of all detected proteins (n=4,705) across three experimental comparisons: AD+P vs NC (left), AD-P vs NC (middle), and AD vs NC (right).

**Supplementary Figure 2.**
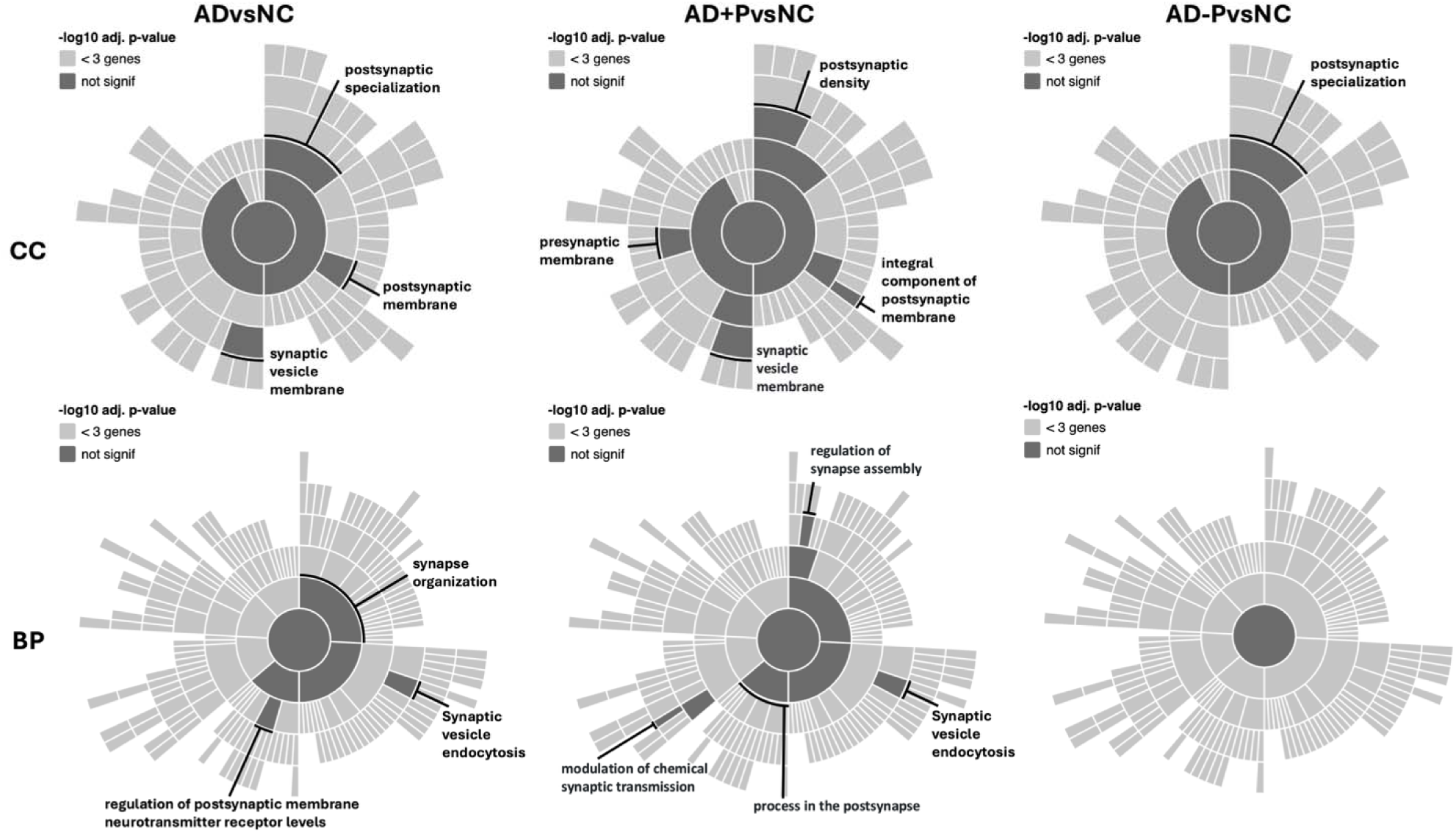
SynGO enrichment analysis. Sunburst plots showing SynGO cellular component (top row) and biological process (bottom row) enrichment analyses for three comparisons.

To explore whether the diverse molecular disruptions in AD+P reflect altered gene co-expression patterns, we applied weighted gene co-expression network analysis (WGCNA)^26^. WGCNA identified 15 distinct co-expression modules, ranging from 50 genes (lightcyan) to 1,588 genes (turquoise) (Fig. 3A, Supplementary Data 6). Module trait relationship analysis revealed four modules (magenta, purple, midnightblue, and tan) that were significantly dysregulated across all comparisons (q < 0.05 for AD vs NC, AD+P vs NC, and AD-P vs NC) (Fig. 3B, 3C, Supplementary Data 7). Notably, these four modules showed similar dysregulation patterns regardless of psychosis status, suggesting they represent core molecular changes in Alzheimer’s disease pathology.

**Figure 3.**
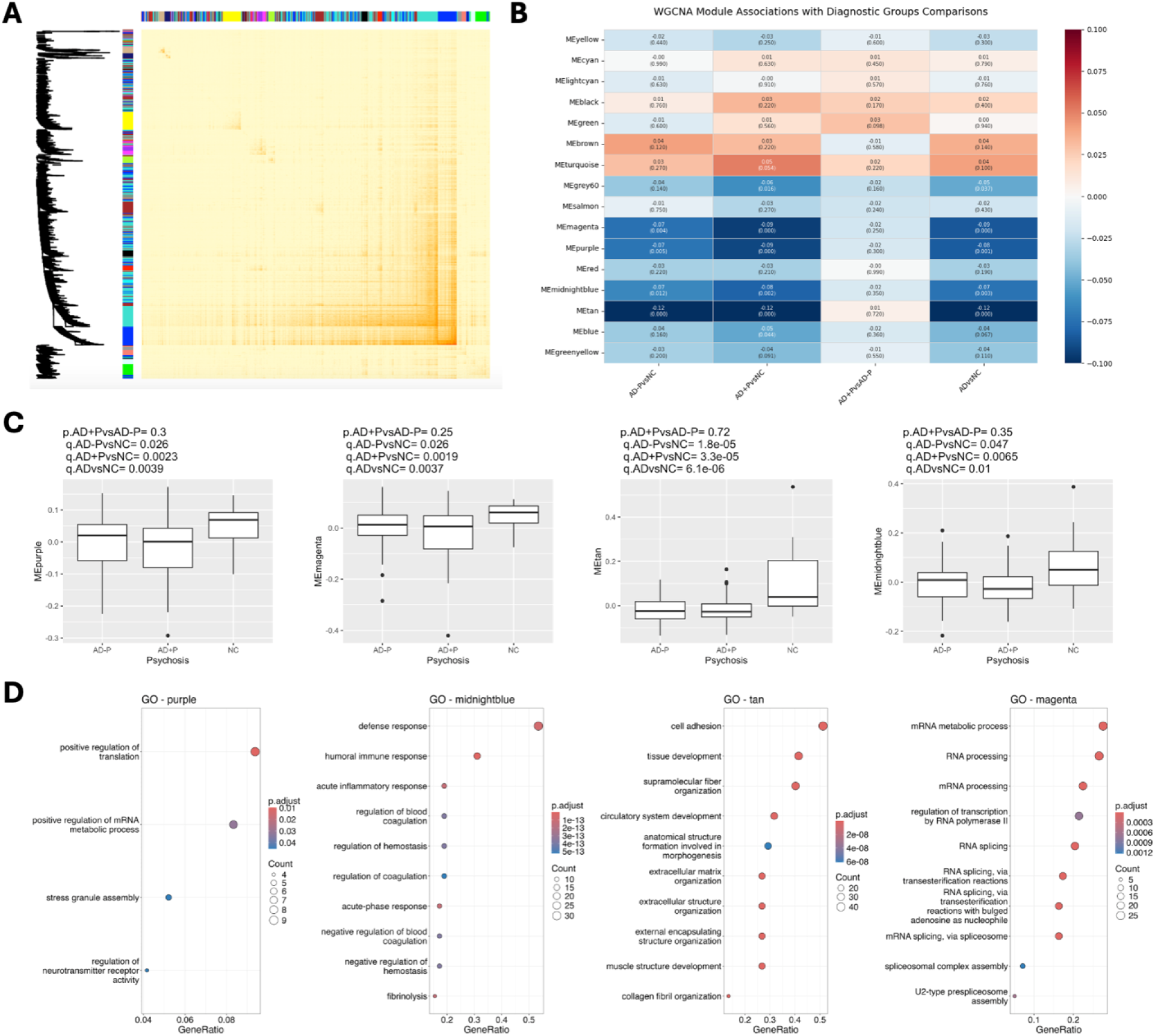
Weighted Gene Co-expression Network Analysis (WGCNA). **A.** Heatmap representing topological overlap matrix with corresponding hierarchical clustering dendrogram and module color of proteins. **B.** Heatmap showing Pearson correlations between module eigengenes and clinical traits, including AD with psychosis (AD+P), AD without psychosis (AD–P), and non-demented controls (NC). Each cell contains the correlation coefficient (r) and p-value; red indicates positive and blue indicates negative correlations, with color intensity reflecting magnitude. **C.** Boxplots showing eigengene expression values for four representative modules across psychosis groups (AD+P, AD–P, NC). Boxes represent the interquartile range (IQR; 25th–75th percentile), the center line marks the median, and whiskers extend to 1.5× IQR. Individual points indicate outliers. p-values for pairwise comparisons are provided above each plot. **D.** Bubble plots of Gene Ontology (GO) biological process enrichment for each module. Bubble size represents the number of proteins annotated to each term, and color indicates the adjusted p-value (FDR).

Functional enrichment analysis of the four core dysregulated modules revealed distinct biological processes (Figure3D, Supplementary Data 8-11). The tan module was enriched for cell adhesion processes, the purple module was enriched for positive regulation of translation, the midnightblue was enriched for defense response, and the magenta module was enriched for mRNA metabolic processes. This WGCNA further supports the previous conclusion by demonstrating that core AD-related disruption occurs similarly regardless of psychosis status.

### Proteins correlated with PSD yield

We next evaluated the relationship of homogenate protein abundances with PSD yield. Out of 4,705 quantified proteins, 15 proteins showed significant correlations with PSD yield (q < 0.05), with eight positively correlated (TBCB, DAAM2, APOB, RABGGTB, IQGAP1, ENPP6, CDH20, and KNG1) and seven negatively correlated (PCSK1, CRACDL, SH3GL2, SLC6A17, GABRG2, SEMA4D, and SURF4). An additional 108 proteins had correlations with q values < 0.1) (Fig. 4, Supplementary Figure 4, Supplementary Data 12). To assess whether these associations were independent of demographic and pathological covariates, we performed multiple linear regression adjusting for sex, age, PMI, APOE4 status, amyloid-beta volume fraction, and tau volume fraction. The covariate-adjusted regression coefficients were highly correlated with the Pearson correlation coefficients (R² = 0.874; Supplementary Figure 3), suggesting that the correlations of homogenate proteins with PSD yield were in large extent not accounted for by AD pathologic burden.

**Figure 4.**
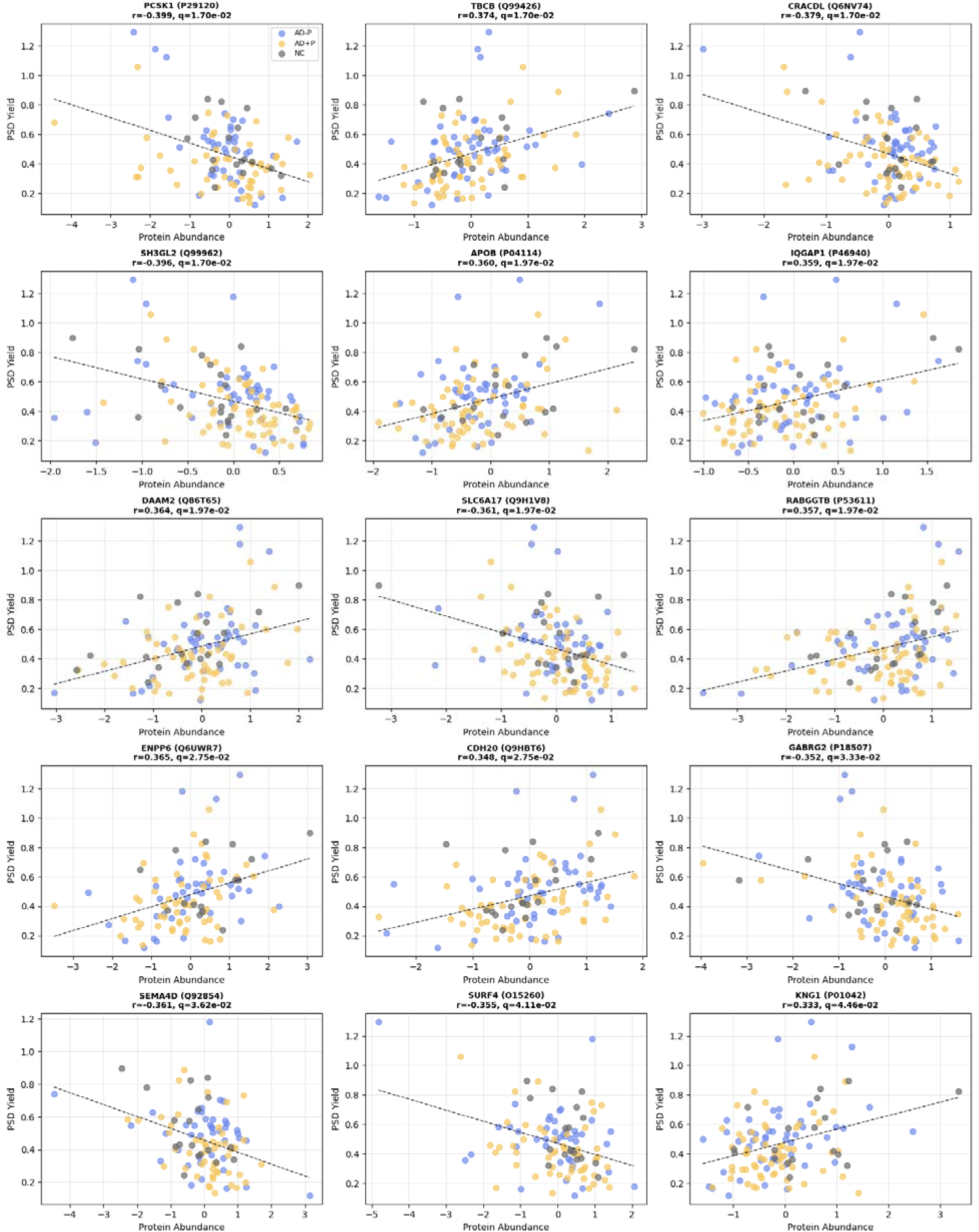
Correlation between protein abundance and PSD levels across diagnostic groups. Scatter plots show the relationship between the abundance of selected proteins and PSD levels in the dorsolateral prefrontal cortex across normal controls (NC, gray), Alzheimer’s disease without psychosis (AD-P, blue), and Alzheimer’s disease with psychosis (AD+P, yellow). Each panel represents one protein significantly correlated with PSD (q < 0.05). Pearson’s correlation coefficient (r) and false discovery rate-adjusted p-value (q-value) are indicated. Dashed lines represent the linear regression fit across all samples.

To complement our individual protein analysis and identify coordinated biological pathways associated with PSD yield, our WGCNA analysis revealed 1 module (midnight blue) significantly correlated with PSD yield across all samples, and 5 modules that had correlations with q values < 0.1) (Fig. 5, Supplementary Figure 5, Supplementary Data 13).

**Figure 5.**
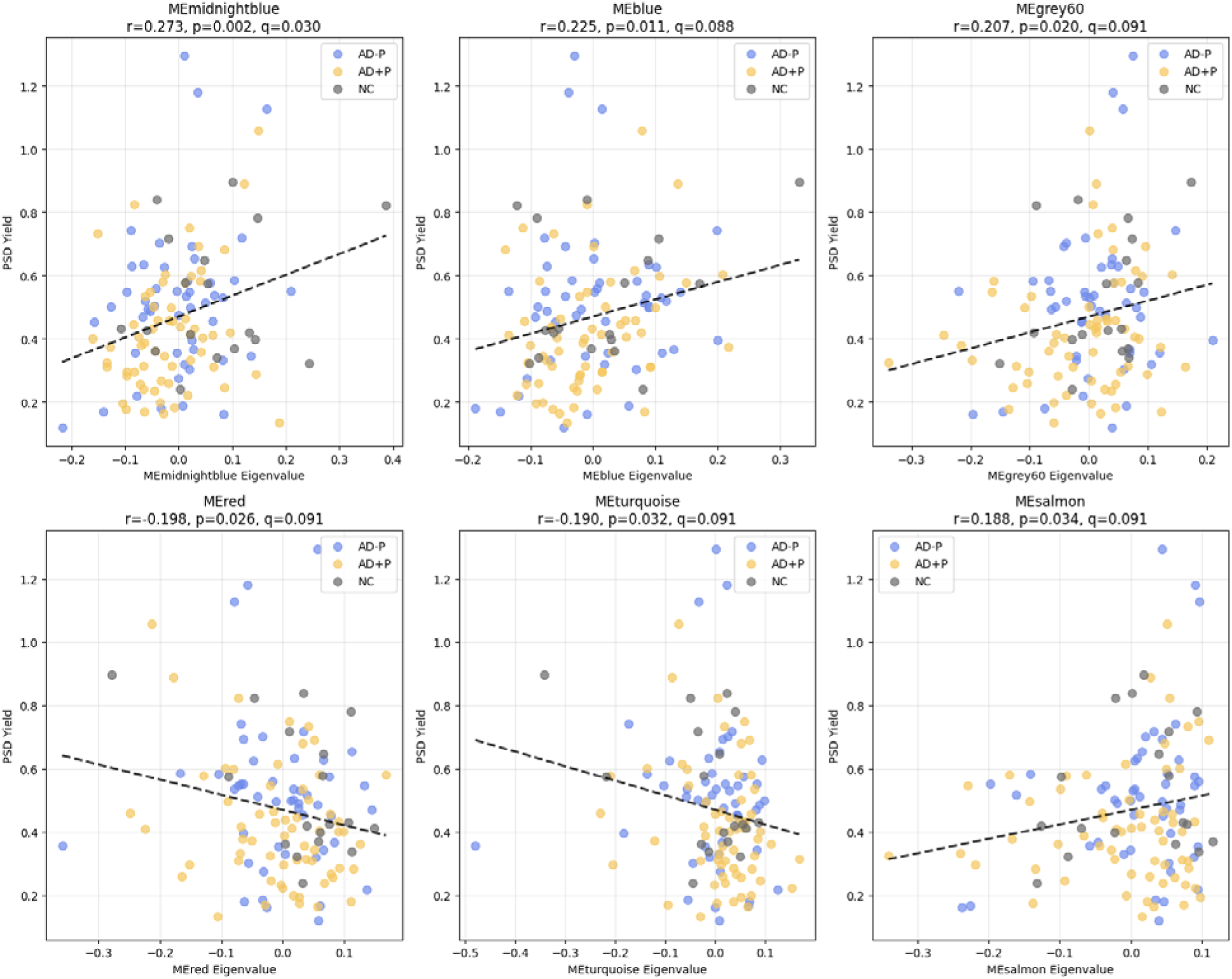
Correlations between module eigengenes and postsynaptic density (PSD) levels. Scatter plots showing relationships between selected module eigengenes (MEs) and PSD levels across diagnostic groups: Alzheimer’s disease with psychosis (AD+P, yellow), Alzheimer’s disease without psychosis (AD-P, blue), and normal controls (NC, gray). Each point represents an individual sample. Pearson’s correlation coefficient (r), p-value, and false discovery rate-adjusted q-value are shown. Dashed lines indicate linear regression fits across all samples.

To determine whether PSD yield-correlated proteins were a subset of the broad AD proteomic signature, we then examined the overlap between the two sets. Proteins most strongly correlated with PSD yield showed only one protein (APOB) overlapped with those altered in AD.

To identify coordinated pathway-level alterations associated with PSD abundance independent of differential expression thresholds, we performed GSEA on proteins ranked by correlation with PSD abundance (Supplementary Figure 6, Supplementary Figures 14, 15). GSEA identified significant enrichment of pathways related to coagulation, steroid metabolic process, and wound healing among proteins positively correlated with PSD abundance. In contrast, proteins negatively correlated with PSD abundance were enriched for pathways related to presynaptic endocytosis, synaptic vesicle recycling, vesicle-mediated transport, and clathrin-dependent endocytosis. Together, these findings indicate that PSD yield-related proteins may capture a distinct biological process from those broadly disrupted in AD, and that APOB may represent a critical link between general AD pathology and psychosis-specific mechanisms.

**Supplementary Figure 3.**
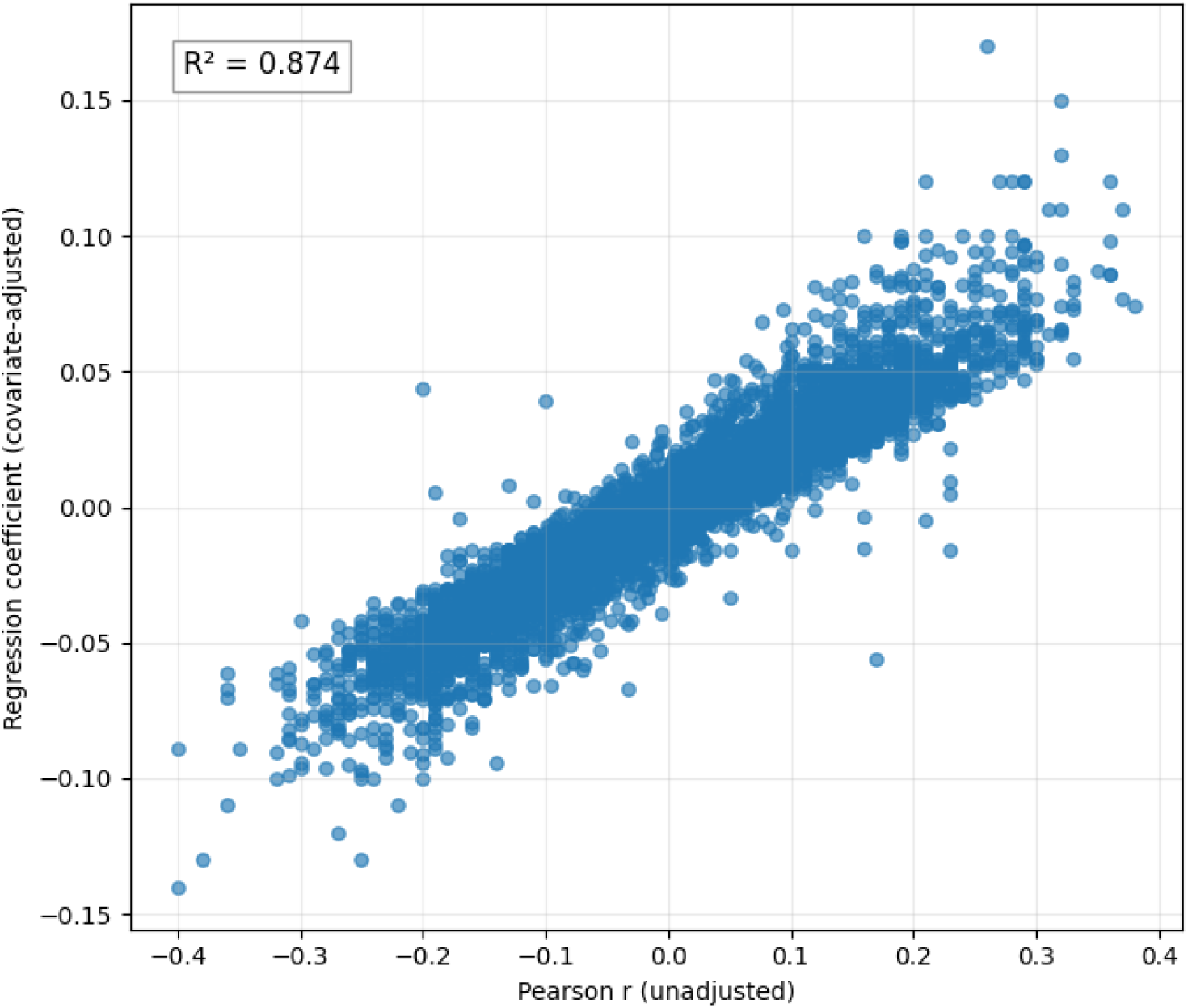
Concordance between Pearson correlation coefficients and covariate-adjusted regression coefficients for PSD yield-associated proteins. Scatter plot comparing Pearson correlation coefficients (x-axis) and covariate-adjusted regression coefficients (y-axis) for all quantified proteins. Covariates included sex, age, PMI, APOE4 status, amyloid-beta volume fraction, and tau volume fraction. The two measures were highly correlated (R² = 0.874, p<2.2e-308).

**Supplementary Figure 4.**
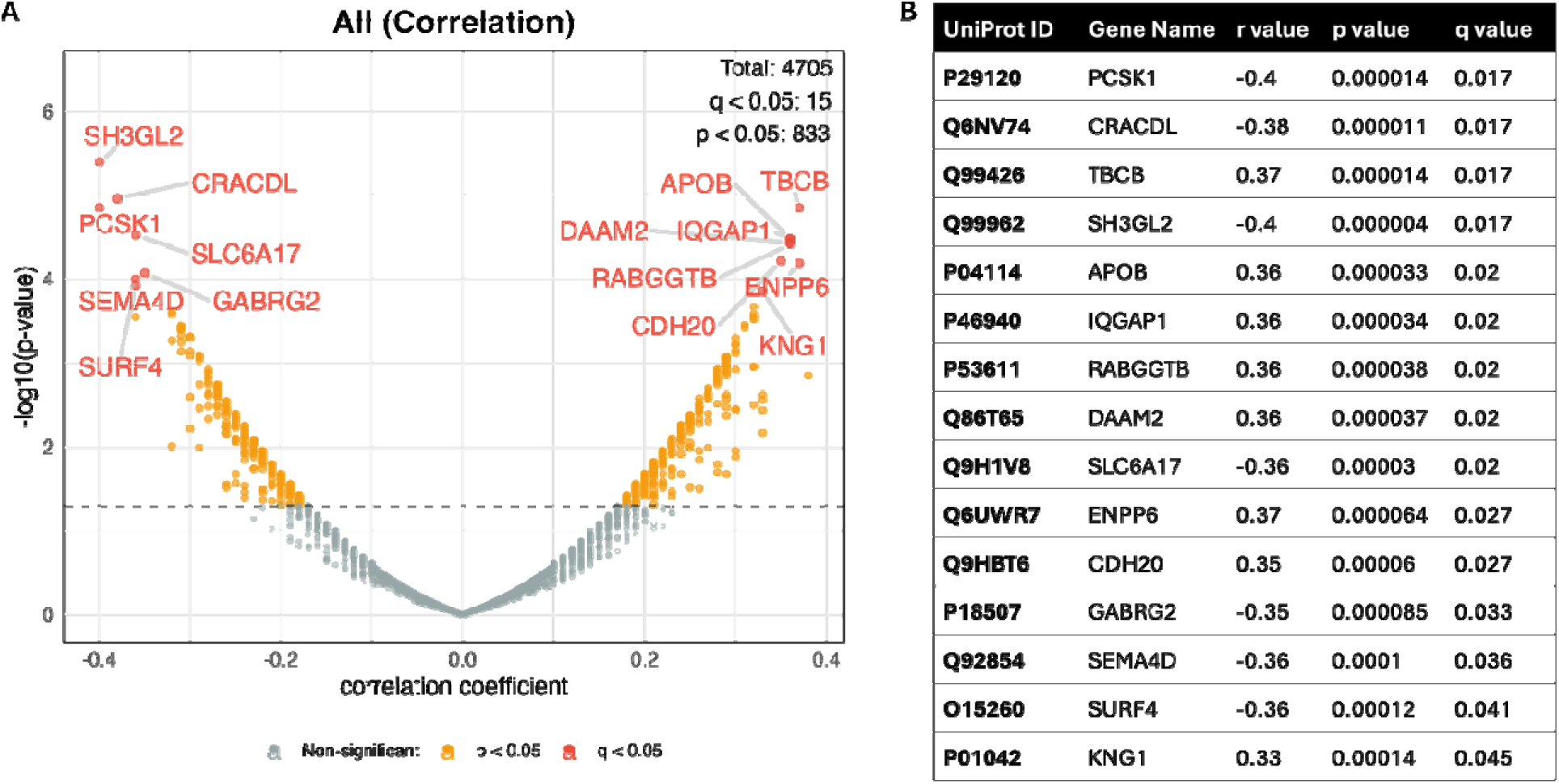
Correlation between protein abundance and PSD yield. (A) Computed Pearson’s correlation between each protein’s abundance and the PSD yield. The dashed line indicates p = 0.05. Red dots and orange dots represent proteins with q < 0.05 and p < 0.05, respectively. (B) The top 15 proteins most positively and negatively correlated with PSD yield (by q value) are labeled and listed in the accompanying table with their UniProt ID, gene name, r value, p value, and q value.

**Supplementary Figure 5.**
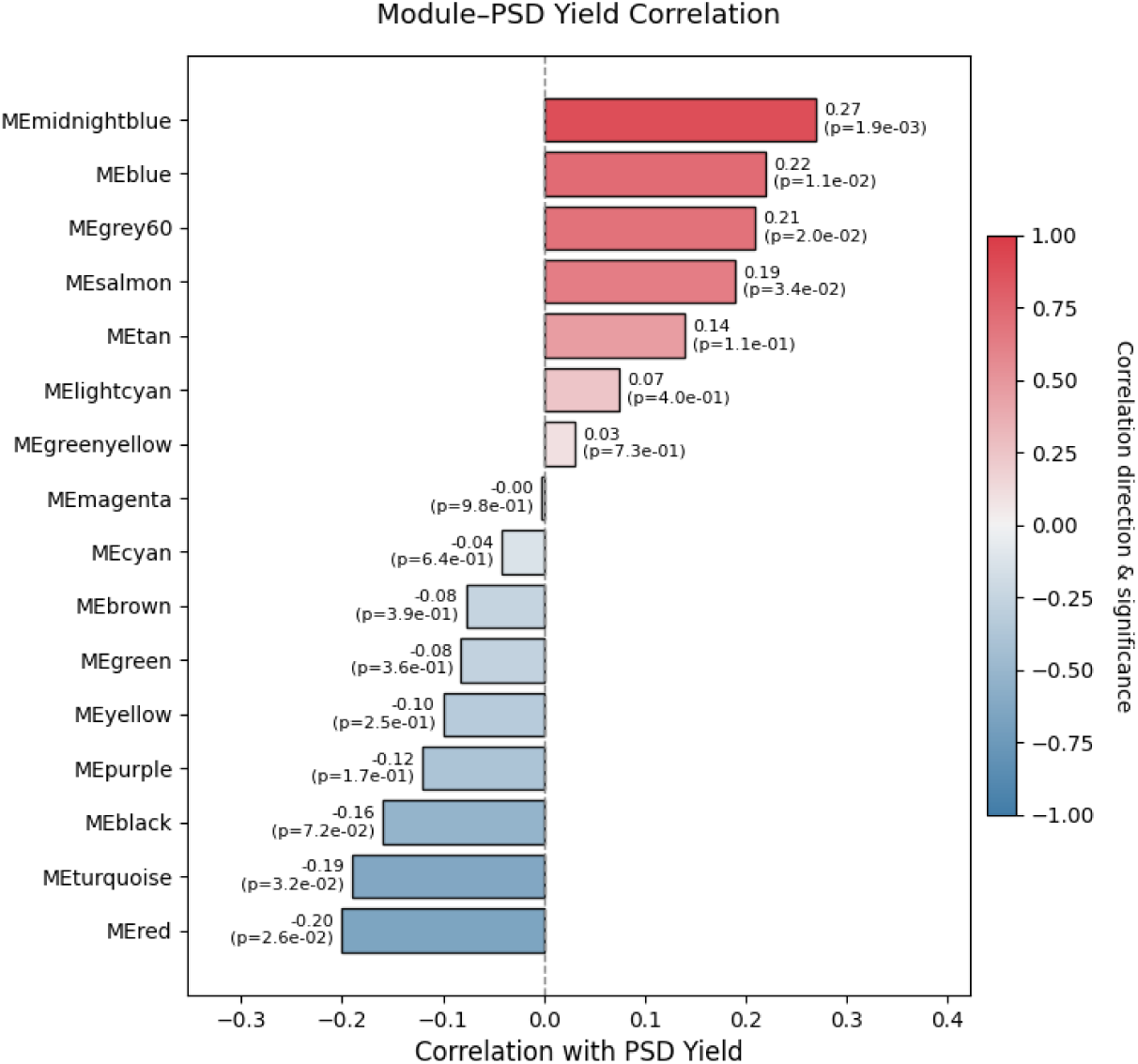
Correlation between module eigengenes and PSD levels. Horizontal bar plot displaying Pearson correlations between module eigengenes (MEs) and PSD levels across all samples. Each cell shows the correlation coefficient (top) and corresponding p-value (bottom). Red indicates positive correlations, and blue indicates negative correlations.

**Supplementary Figure 6.**
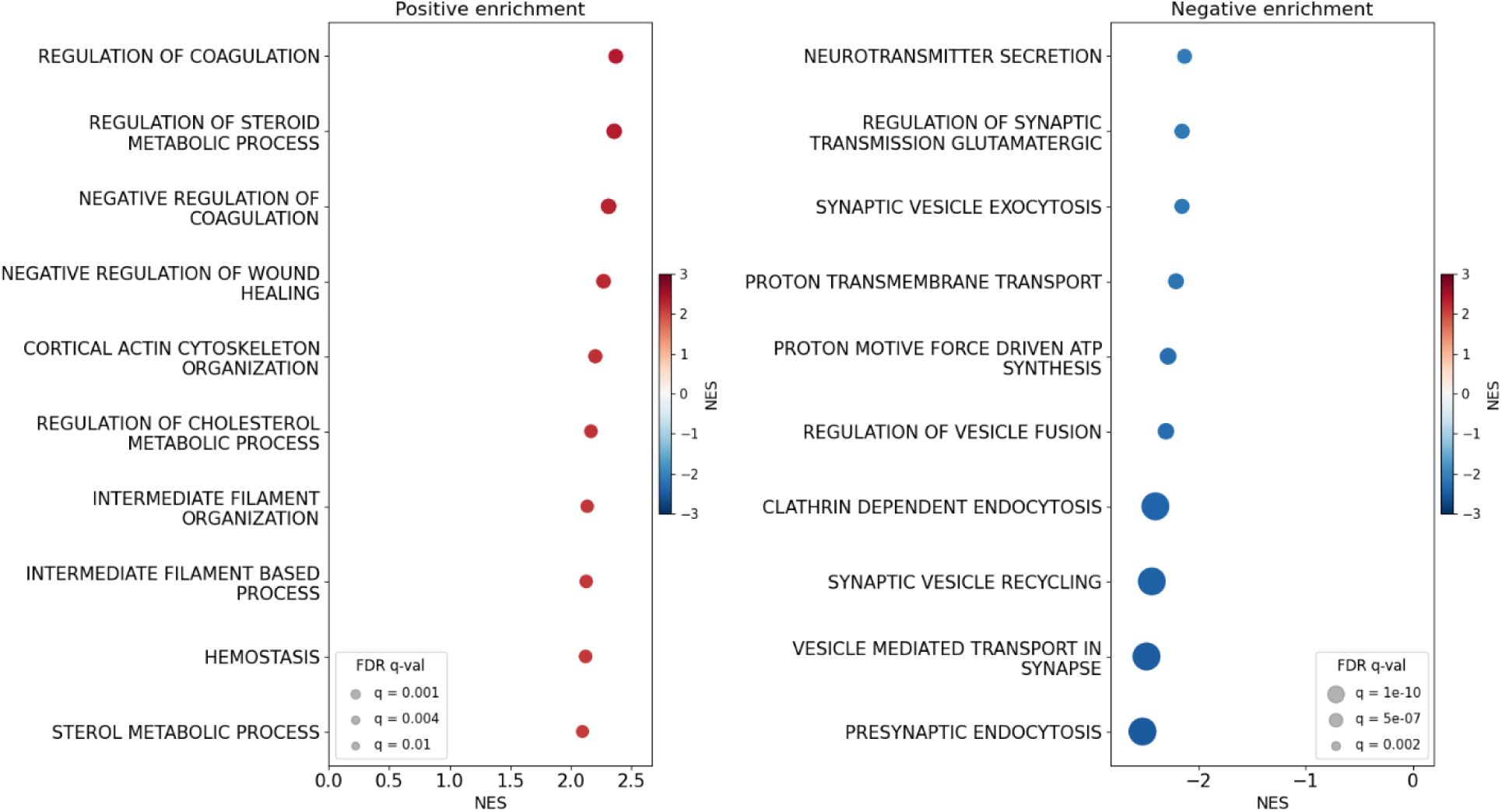
Bubble plot of GSEA results for proteins positively (left) and negatively (right) correlated with PSD yield. The top 10 gene sets with FDR q < 0.25 are shown. Bubble size reflects statistical significance (−log₁₀ FDR q-value); color and x-axis position indicate normalized enrichment score (NES).

## Discussion

In this study, we profiled dorsolateral prefrontal cortex gray matter proteomes from individuals with Alzheimer’s disease with and without psychosis to delineate molecular alterations underlying psychotic symptoms. We found that AD+P exhibits the greatest number of differentially expressed proteins relative to controls, yet proteomic changes were highly correlated between AD+P and AD–P, suggesting that psychosis does not arise from a distinct set of proteomic alterations in gray matter as a whole, but rather reflects quantitative shifts within a shared AD proteomic landscape. These results of gray matter proteomes were striking in their difference from our prior study of the PSD proteome in these same subjects, in which AD+P showed substantial reductions in PSD yield and PSD protein abundances relative to AD-P and to elderly control subjects. Examination of gray matter homogenate protein correlations with PSD yield revealed them to be largely independent of the broad AD proteomic signature and of AD proteinopathy, suggesting that PSD-related molecular alterations may capture unique mechanisms of synaptic vulnerability linked to psychosis risk in AD.

The strong correlation between AD+P and AD–P gray matter homogenate proteomic profiles and overlapping enrichment of extracellular matrix and structural organization pathways support the idea that psychosis occurs within a background of a common AD molecular framework. Prior studies similarly reported that AD+P and AD–P show comparable amyloid, and only modestly different tau burdens, with psychosis reflecting additional network-level or synaptic dysfunction rather than divergent pathology^5,15^. The enrichment of supramolecular fiber and extracellular matrix organization pathways across all comparisons suggests a robust signature of structural remodeling in the AD cortex, consistent with recent proteomic datasets highlighting ECM dysregulation and glia activation as central features of AD progression^27,28^. These shared processes likely reflect broad synaptic and vascular alterations, not specific to psychosis.

Proteins correlated with PSD yield provide an independent perspective on mechanisms of synaptic vulnerability. Several proteins positively correlated with PSD represent protective synaptic factors whose loss-of-function may contribute to psychosis vulnerability. TBCB (Tubulin-folding cofactor B) regulates neuronal axonal growth and is down-regulated in schizophrenia cell lines^29^ and negatively correlated with AD pathology^30^, suggesting its reduction may be associated with compromised synaptic integrity. IQGAP1 plays a critical role in NMDA receptor trafficking and synaptic plasticity^31^, with knockout studies demonstrating severe long-term memory deficits and impaired hippocampal LTP^32^. ENPP6, a choline-specific glycerophosphodiesterase that is essential for oligodendrocyte maturation^33^, is especially notable as it was identified as a risk factor for AD+P in a previous GWAS study^34^. Genetic variants in ENPP6 increase susceptibility to AD+P, and our proteomic finding that higher ENPP6 levels correlate with preserved PSD yield raise a potential loss-of-function mechanism, though the precise molecular pathways remain to be established.

APOB presents a notable case as the only protein among those most strongly correlated with PSD yield that was also significantly altered in AD. APOB was reduced in both AD+P and AD-P, indicating its involvement in general AD pathology. APOB has been linked to AD and dementia risk^35,36^, shows enrichment of rare coding variants in early-onset AD cases^37^, and has elevated cerebrospinal fluid levels associated with tau pathology in presymptomatic individuals^38^. Previous work found that APOB protein levels in the prefrontal cortex were positively correlated, whereas APOB mRNA levels in the DLPFC were negatively correlated, with AD pathology and cognitive impairment ^39^. The latter finding is directionally consistent with our observation of reduced APOB protein levels in AD, whereas the protein-level findings contrast with our results. These discrepancies highlight the complexity of APOB regulation across molecular levels. Although APOB has been implicated in AD-related genetic risk, how these associations manifest at the protein level remains unclear. Further studies will be needed to determine whether APOB represents a reproducible biological association or a false-positive finding.

In contrast, proteins negatively correlated with PSD yield show distinct relationships to AD pathology. SEMA4D is up-regulated in AD and has been shown to directly impair synaptic function by binding to astrocytes and reducing EAAT-2 glutamate transporter and glutamine synthetase level^40^. A SEMA4D-blocking antibody (pepinemab) is currently in clinical trials for neurodegenerative diseases. SURF4 has been reported to increase the neurotoxicity of Aβ1-42 peptides^41^ and serves as a cargo receptor for APOB secretion^42^. The negative correlation of these proteins with PSD yield is consistent with their documented roles in contributing to synaptic dysfunction in AD, though their specific relevance to AD+P remains to be established. SH3GL2, involved in synaptic vesicle endocytosis, is also up-regulated in AD^43^ and has been identified as a shared gene in AD and diabetes-associated cognitive dysfunction^44^.

Further insight may be gleaned from the processes enriched for proteins positively and negatively correlated with PSD yield. Although the positively enriched pathways were annotated primarily as coagulation-, wound-healing-, and steroid metabolism-related processes, several leading-edge proteins within these pathways have established roles in neuronal signaling and synaptic maintenance. EPHB2, a key contributor to these pathways, is a well-established regulator of excitatory synapse development^45,46^, dendritic spine morphology^47^, and restoration of EPHB2 expression rescues synaptic and cognitive deficits in AD models^48,49^. Other leading-edge proteins, including VTN and KNG1, have been linked to neurite growth, integrin signaling^50^, and CDC42-dependent cytoskeletal remodeling^51^, processes important for synaptic plasticity.

The enrichment of synaptic vesicle recycling, presynaptic endocytosis, and vesicle trafficking pathways among proteins negatively associated with PSD suggests that reduced postsynaptic integrity is accompanied by increased engagement of presynaptic machinery. One potential interpretation is that these changes reflect compensatory presynaptic remodeling in response to postsynaptic dysfunction. Experimental studies have demonstrated that reduced postsynaptic activity can trigger retrograde signaling mechanisms that enhance presynaptic release probability and vesicle cycling through pathways involving BDNF-TrkB signaling^52^, and mTORC1-dependent translation^53^. Thus, the enrichment of presynaptic trafficking pathways among proteins negatively correlated with PSD yield may represent an adaptive attempt to maintain synaptic transmission despite postsynaptic deficits.

This latter interpretation, i.e. of a primary postsynaptic deficit in AD+P, aligns with our study of the PSD proteome in AD+P^7^. That study revealed enrichment for reduced levels of a network of 23 kinases and proteins regulating the postsynaptic actin cytoskeleton. This network, which is essential to synapse formation and maintenance^54^, is one in which protein levels may be less relevant than their activities, which are largely controlled by phosphorylation.

There is, however, a discordance between the observed reduction in PSD yield in AD+P and the absence of significant individual protein differences between AD+P and AD-P in bulk brain homogenate. The majority of PSDs in cortex are found in dendritic spines of excitatory neurons, thus, the reductions in DLPFC PSD yield in AD+P are likely to reflect selective excitatory neuron vulnerability^9,55^. The bulk gray matter analysis in the current study may have masked detection of protein-level alterations due to signal dilution across heterogeneous cell types and subcellular compartments.

Despite the limitations in identifying specific proteomic differences between AD+P and AD-P, these findings highlight candidate pathways of potential translational relevance in a hypothesis-generating manner. The convergence of genetic and proteomic evidence for ENPP6 provides the strongest rationale for therapeutic intervention. ENPP6’s role in generating sphingosine creates a direct mechanistic link to fingolimod, an FDA-approved sphingosine-1-phosphate receptor modulator. Preclinical studies have demonstrated that fingolimod rescues psychosis-associated behavioral deficits and increases synaptic protein abundance in AD mouse models^56^. Additionally, SEMA4D represents another actionable target, as pepinemab, a SEMA4D-blocking antibody currently in clinical trials for neurodegenerative diseases, may benefit synaptic preservation through its effects on astrocytic glutamate homeostasis (Reviewed in^57^). Together, these observations nominate ENPP6- and SEMA4D-related pathways as candidates for further investigations.

Collectively, our findings suggest a model in which psychosis in Alzheimer’s disease arises from the combination of quantitative alterations within a shared AD proteome profile and a superimposed set of protein alterations correlated with PSD yield that are largely independent of the shared AD proteome, are independent of AD proteinopathy burden, and which may confer synaptic vulnerability. Future studies of AD+P may benefit from examining phosphoproteomic alterations within PSD and directly examining the presynaptic compartment’s proteome and phosphoproteome to uncover additional targets for intervention.

## Acknowledgments

This work was supported by the National Institutes of Health (R01 MH116046, P30 AG066468, R01MH125235 and R01MH118497).

